# Cyclic voltammetry electropolymerization of PEDOT:Polydopamine on gold electrodes^*^

**DOI:** 10.64898/2026.05.27.728128

**Authors:** Saloua Saghir, Giuseppe Schiavone

## Abstract

This work presents a facile and rapid fabrication method for high-performance bioelectrode coatings through electropolymerization of poly(3,4-ethylenedioxythiophene) (PEDOT) and polydopamine (PDA). Expanding on previous work, we develop a cyclic voltammetry electropolymerization process to deposit PEDOT:PDA coatings on 2 mm gold electrode substrates. The coatings exhibit a 40-fold increase in cathodic charge storage capacity (~ 40mC·cm^−2^) and significant impedance modulus reduction compared to uncoated gold electrodes. Morphological characterization revealed a uniformly porous surface that corroborates the enhancement in electrochemical performance. Our scalable approach offers a promising option to fabricate bioelectrodes for application to neural interfaces and implantable and wearable bioelectronics.

**Clinical Relevance:** The improved electrochemical performance and scalable fabrication of PEDOT:PDA coatings support their potential to enhance the stability and signal quality of neural and implantable bioelectronic devices.

## I. Introduction

Conductive polymers (CP) have spiked great interest in recent years owing to their unique combination of properties such as tunable mechanical signature, dual electronic-ionic conduction, processability, and possible bio-chemical functionalization. Applications include flexible electronics, energy storage, electrochemical sensors, neuroelectronic interfaces, wearable bioelectronics, and tissue engineering scaffolds [1]. Among popular CPs, poly(3,4-ethylenedioxythiophene) (PEDOT) stands out for its high electrical conductivity and biocompatibility. In addition, PEDOT can be functionalized with biomolecules such as polydopamine (PDA) to enhance biointerface performance by leveraging PDA’s properties such as adhesiveness, antibiofouling capability, biocompatibility and its numerous functional groups for chemical functionalization. In this work, we report a method for the electropolymerization of PEDOT and PDA producing low-impedance, high charge storage capacity coatings on 2 mm diameter gold electrodes.

While we have previously reported on the synthesis and characterization of PEDOT:PDA coatings by potentiostatic deposition (PS) [2], here we demonstrate that cyclic voltammetry (CV) can equally be employed as an alternative electrodeposition technique, achieving comparable results and therefore expanding the manufacturing compatibility of PEDOT:PDA to fit needs such as available instrumentation, user preferences, or specific experimental workflows. Providing researchers with multiple fabrication options facilitates wider adoption of PEDOT:PDA coatings and supports method optimization for diverse bioelectronic applications.

## II. Methods

### A. Fabrication of the electrode substrates

Electrode substrates were fabricated according to a previous protocol [2], using a rapid prototyping method. Briefly, four-inch silicon wafers are thermally oxidized to form a 200 nm silicon dioxide layer and subsequently coated with 10 nm of chromium and 100 nm of gold via magnetron sputtering at 0.1 Å/s and 0.65 Å/s rate, respectively. A 65 µm thick Kapton tape passivation is then laminated onto the metallized wafer and then patterned by CO_2_ laser to expose 2 mm diameter circular gold electrodes, and contact pads for electrical connection to a potentiostat (CH Instrument CHI660E). Wafer dicing using a diamond saw is used to singulate individual chips comprising of an electrode and a connection pad. The cross-sectional edges of the chip where the thickness of the sputtered metals is exposed is then passivated with silicone all around the die, to prevent Kapton tape delamination and to ensure that electrical conduction in the solution is confined to the working electrode.

### B. Electropolymerization of PEDOT:PDA

Electrodeposition is carried out in a three-electrode setup, with the fabricated gold electrode as the working electrode, a Ag/AgCl reference electrode, and a large-area platinum mesh counter electrode. The electrodeposition solution is prepared by separately dissolving i) 10 mg of dopamine in 9 mL phosphate buffer saline (PBS) at pH 7.2; and ii) 10.7 µL EDOT (10 mM) in 1 mL ethylene glycol. Next, the two solutions are mixed under magnetic stirring for 3 min and purged under nitrogen for 10 min. CV deposition is carried out at 50 mV/s scan rate, with a voltage range of −1 V to +1.2 V. The deposition is stopped once 50 mC deposition charge is reached at +1 V. Fig. 1 illustrates the fabrication process of the coated electrodes.

**Figure 1:**
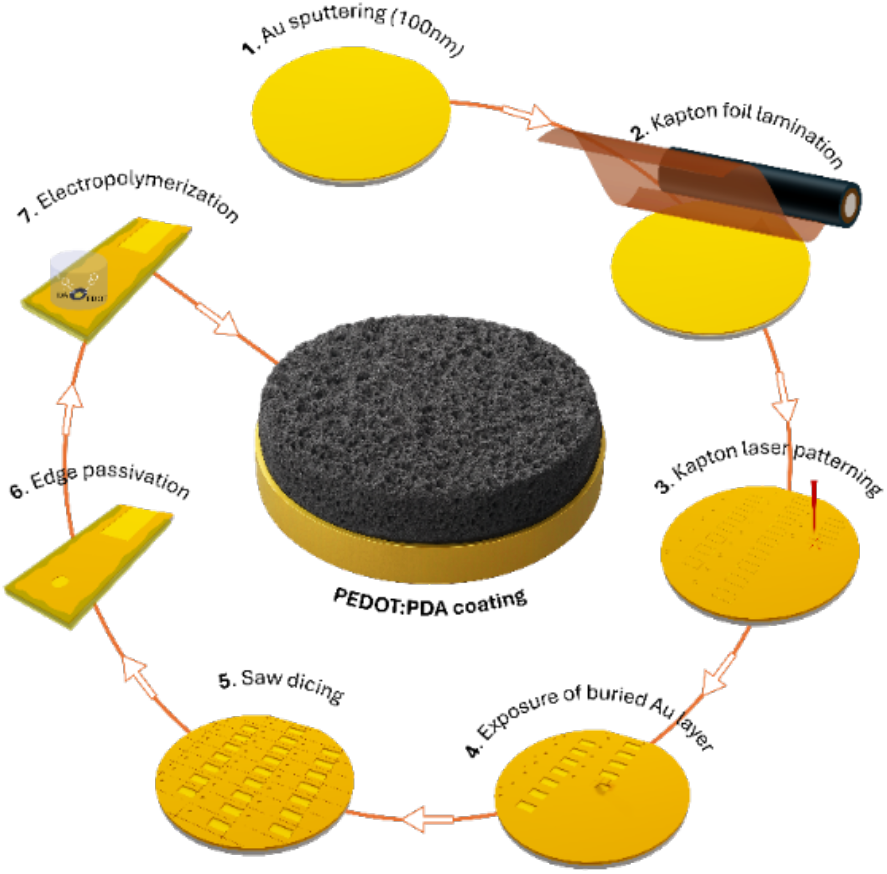
Fabrication process of PEDOT:PDA coated gold electrodes.

### C. Electrochemical characterization

After electropolymerization, the coated electrodes were characterized by electrochemical impedance spectroscopy (EIS) and CV in a standard three-electrode configuration [3].

EIS measurements were conducted in PBS with 10 mV amplitude, 0 V DC offset vs. open circuit potential, and a frequency sweep from 1 Hz to 100 kHz. CV measurements were performed in 0.1 M KCl containing 2 mM potassium hexacyanoferrate(III) with voltage sweep from −0.6 V to 0.8 V at a 100 mV/s scan rate. Five consecutive CV scans are performed prior to each recorded measurement, to stabilize the voltametric response.

### D. Morphological and topographical characterization

The PEDOT:PDA thin films were also characterized via scanning electron microscopy (SEM Hitachi SU 8230) and atomic force microscopy (AFM XE-200, Park Systems), to respectively evaluate topography and roughness on a 25 µm by 25 µm scan area. The coating thickness was characterized using a mechanical profilometer (Bruker Dektak) after removing the Kapton insulation. Each sample was scanned at 3 positions along the coated area (top, middle, and bottom).

### E. Structural characterization

ATR-FTIR spectroscopy (Nicolet iS50, Thermo Fischer) was used to characterize the chemical composition of pure PEDOT, pure PDA, and the PEDOT:PDA composite coating. Pure PEDOT powder was obtained by self-polymerization of EDOT following a previously described protocol [4]. Briefly, 114 µL of EDOT was dissolved in 4 mL ethanol, to which 9.4 mL of formic acid was added. The solution was kept in a sealed vessel to avoid solvent evaporation and left to polymerize for 3 weeks at room temperature. The resulting PEDOT solution was poured into a Petri dish and left to dry at room temperature until complete solvent evaporation. The dried PEDOT film was then rinsed with ethanol, scraped from the glass surface, and the collected powder was left to dry prior to analysis. Pure PDA powder was prepared by dissolving 500 mg dopamine in 3 mL 1 M NaOH. The solution was poured into a Petri dish and placed in an oven at 40°C until complete solvent evaporation to obtain PDA powder. For the PEDOT:PDA composite, the coating was deposited as per the method described in section II.B onto a gold-sputtered Kapton sheet to ensure good conformable contact with the ATR crystal during measurement.

### F. Error calculations

Measurements are reported as mean ± standard error (SE), calculated as

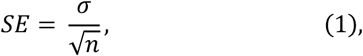

where σ is the standard deviation and *n* the number of measurement samples.

## III. Results

### A. PEDOT:PDA electropolymerization

CV is an electropolymerization technique in which a potential is swept across a specified voltage range, and the resulting current is recorded. The number of cycles will define the thickness of the coating and subsequently the electrochemical performances. The electropolymerization of dopamine is a complex, multi-step mechanism involving multiple intermediate species that can undergo redox reaction as illustrated in Fig. 2 and observed in the CV deposition curves of Fig. 3.

**Figure 2:**
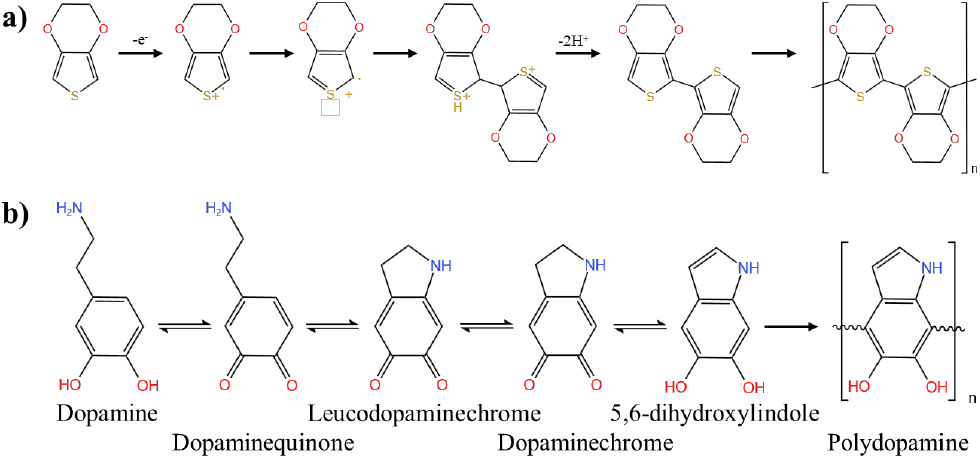
Electropolymerization mechanism of (a) PEDOT and (b) PDA

**Figure 3:**
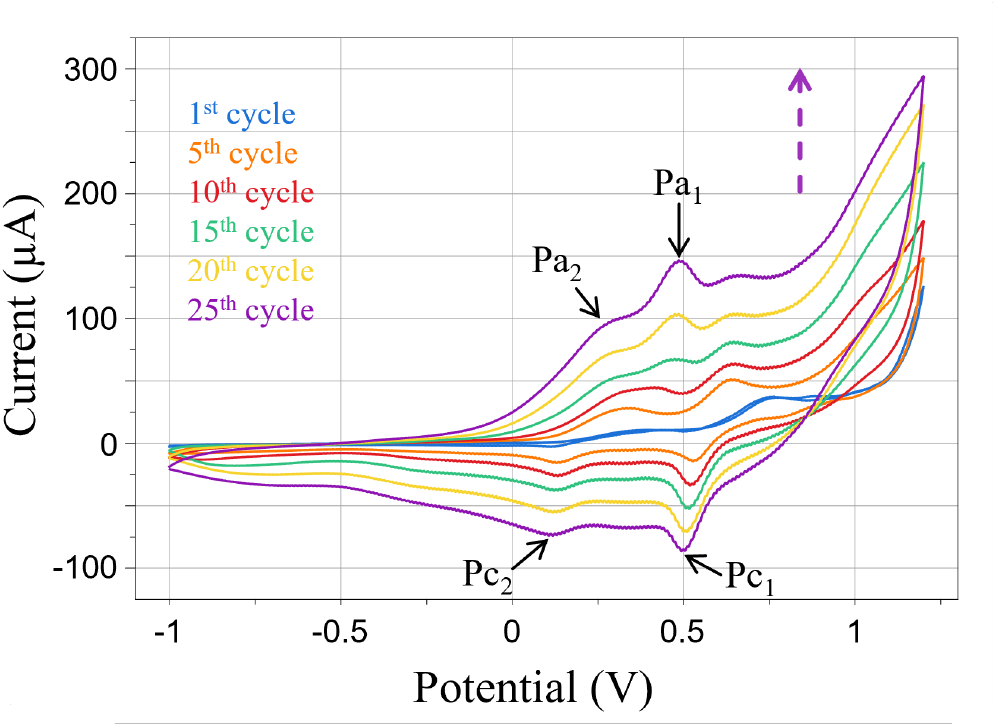
Current-voltage CV deposition curves, between –1.0 V and +1.2 V at a scan rate of 50 mV s^-1^, 25 cycles.

In the CV scans, an anodic peak is observed at +0.5 V and a cathodic peak at −0.5 V, which can be attributed to the dopamine/dopamine quinone redox couple (Pa_1_/Pc_1_). The secondary anodic peak at +0.3 V and cathodic peak at −0.3 V can be attributed to the leucodopaminechrome/dopaminechrome couple (Pa_2_/Pc_2_). These peaks are shifted compared to literature values for dopamine electropolymerization alone, where the typical voltammetric responses occur at approximately −0.2 V, +0.21 V, +0.18 V, and −0.3 V for Pa_1_, Pa_2_, Pc_1_, and Pc_2_, respectively [5], [6]. This shift can be explained by the presence of EDOT as a competing species in the electrolyte and the different pH [7], [8]. Although EDOT typically polymerizes at +0.8 V and above, its presence can interfere with dopamine electropolymerization, shifting the redox peaks of dopamine species toward higher anodic potentials and lower cathodic potentials. The broad peak visible between +0.8 V and +1.2 V corresponds to EDOT oxidation and subsequent polymerization. The peak current amplitudes increase progressively with each CV cycle, indicating continuous growth of the electroactive PEDOT:PDA film on the electrode surface.

### B. Electrochemical characterization

After deposition, the coated electrodes were characterized by CV and EIS to extract typical electrode performance metrics used for bioelectrodes such as the cathodic charge storage capacity (CSC_c_), the electroactive surface area (EASA), and the impedance modulus. CSC_c_ is an indicator of the electrode’s ability to reversibly store and deliver charge and is calculated as the area under the negative current curve:

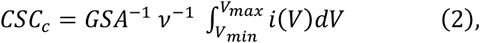

where GSA is the geometric surface area and *v* the scan rate.

The electropolymerized PEDOT:PDA coatings exhibit a significant increase in CSC_c_ to a ~ 40-fold enhancement with respect to uncoated controls (Table 1). Notably, our coating compares favorably to the widely used PEDOT:PSS coatings reported in the field (10 – 35 mC·cm^−2^) [9]. The CV also highlights an increase in electroactive surface area (EASA) to an approximately 9 fold compared to the geometric surface area (0.0314 cm^2^). The EASA is a measure of the actual surface area of the electrode that is electrochemically accessible and participates in charge transduction mechanisms, as opposed to the geometric surface area. The EASA is calculated using the Randles-Sevick equation:

**TABLE 1.** Electrochemical metrics of the bare and pedot:pda coatings.

|  | Gold (n = 3) | CV (n = 3) | PS (n = 22) |
| --- | --- | --- | --- |
| CSC<sub>c</sub><br>(mC·cm<sup>-2</sup>) | 1.08 ± 0.02 | 39.51 ± 4.33 | 42.91 ± 1.74 |
| EASA (cm<sup>2</sup>) | 0.027<br>± 0.001 | 0.27 ± 0.03 | 0.29 ± 0.02 |
| \|z\| at 1Hz (Ω) | 234.30 k<br>± 63.03k | 494.83 ± 59.64 | 379.28 ± 33.78 |
| \|z\| at 1kHz (Ω) | 486 ± 6.18 | 241.53 ± 51.51 | 155.24 ± 4.99 |

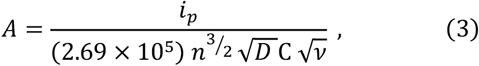

where *A* is the electrochemically active surface area (cm^2^), *i*_*p*_ is the maximum observed peak current in the CV scan, *n* is the number of transferred electrons in the redox reaction (*n* = 1), *D* is the diffusion coefficient of the [Fe(CN)_6_]^3−/4−^ redox probe in KCl solution (7.6 · 10^-6^ cm^2^·s^−1^, [2]), *C* is the concentration of the [Fe(CN)_6_]^3−/4−^ (2 10^−6^ mol·cm^-3^), *v* is the scan rate (V·s^−1^).

Impedance analysis further revealed a significant reduction in impedance modulus, with ~2-fold and ~ 470-fold decrease at 1 kHz and 1 Hz respectively. This is indicative of improved charge transduction with increased interface capacitance, a critical metric for bioelectronic interface applications such as neural recording and stimulation.

### C. Comparison of PS- and CV-deposited PEDOT:PDA

The electrochemical performances of PEDOT:PDA coatings prepared by PS and CV depositions were also compared. To ensure a fair comparison, the literature reports various normalization parameters such as coating thickness, deposition time, or total deposited charge [10],[11]. In this work, we set a fixed deposition charge for the two methods because it provided superior batch-to-batch reproducibility compared to using a fixed number of deposition cycles. The deposition charge calculation differs from a potentiostatic to a potentiodynamic regime. In PS deposition, the current *I* is measured while the potential *E* is held constant over time *τ*, and the total charge *Q* is the direct integration of current *I*:

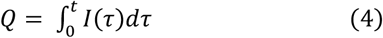

In a potentiodynamic regime such as CV deposition, a potential E is swept linearly at a fixed scan rate *v* and the resulting current is recorded as a function of *E*. Since potential and time are coupled through *v* (equation 5), the deposited charge can be recovered by integrating current over potential rather than time.

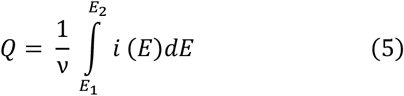

Using these equations, the PS deposition was performed at 1.0 V until reaching a target 50 mC deposition charge, following previously optimized conditions [2]. For CV deposition, the process was stopped after reaching 50 mC when the potential reached 1.0 V during the anodic scan. The number of cycles needed to reach the target 50 mC charge differed across batches, ranging from 24 to cycles. Similarly, the time needed to reach 50 mC in PS deposition differed from sample to sample (600 to 900 s).

Electrochemical performance was assessed using n = 3 bare gold electrodes and n = 3 CV-deposited coatings. For completeness, n = 22 PD-deposited coatings from a previous study were included in the comparison [2]. As displayed in Fig. 4 and reported in Table 1, both coatings offer comparable electrochemical performance across all measured parameters, including CSC_c_, EASA, and impedance modulus at 1Hz and 1kHz. The observed differences fall within the within-group variability, and no statistically significant distinction can be drawn between the two. This demonstrates that PEDOT:PDA coatings can be successfully fabricated through two alternative electrodeposition routes, without compromising electrochemical properties, providing researchers with flexibility in method selection based on available instrumentation and experimental requirements.

**Figure 4:**
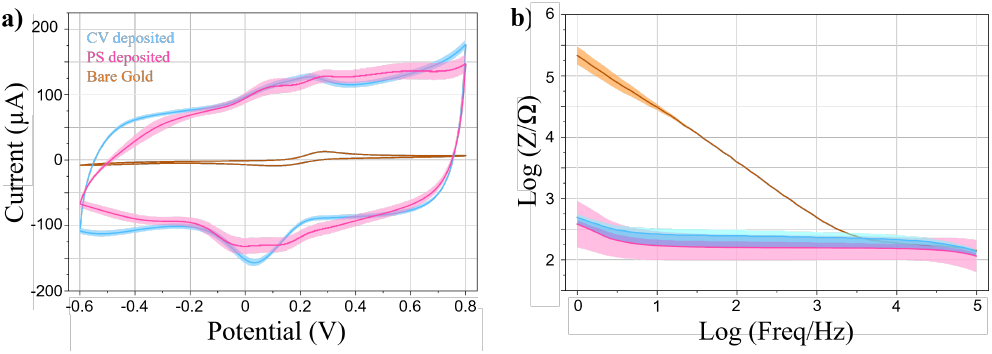
a) CV curves in 2 mM ferro-ferricyanide, 100 mV s^-1^; b) Impedance modulus spectra in PBS. Solid lines: mean values. Shaded areas: SE.

### D. Topography characterization

Optical micrographs (Fig. 5) of the coated electrodes show uniform film coverage for both deposition techniques. SEM images reveal subtle variation in surface topology: PS deposition produced larger, more defined clusters with distinct nodular features and a coarser texture. CV deposition yields finer, more uniformly distributed granular features without clusters distributed across the surface. AFM scans corroborate these observations. PS-deposited coatings show a rougher surface topography, with large, rounded aggregates irregularly distributed across the surface. In contrast, CV-deposited coatings display a more uniform topography, with smaller features evenly distributed across the scan area, resulting in a more homogeneous surface. AFM also reveals very rough surface topologies, with an arithmetic roughness (R_a_) value ~2-fold higher for PS than CV (Table 2). This morphological difference can be attributed to distinct electropolymerization kinetics and growth mechanisms. Potentiostatic deposition reaches a total deposition charge of 50 mC in 10 – 15 min, whereas CV requires longer times (40 – 60 min, depending on the number of cycles until reaching the target charge).

**TABLE 2.** Thickness and surface roughness of measured on pedot:pda samples coated via cv and ps deposition.

|  | CV (n = 3 scans) | PS (n = 3 scans) |
| --- | --- | --- |
| Thickness (nm) | 2915.74 ± 37.86 | 4232.96 ± 10.27 |
| $R_a$ (nm) | 265 | 495 |

**Figure 5:**
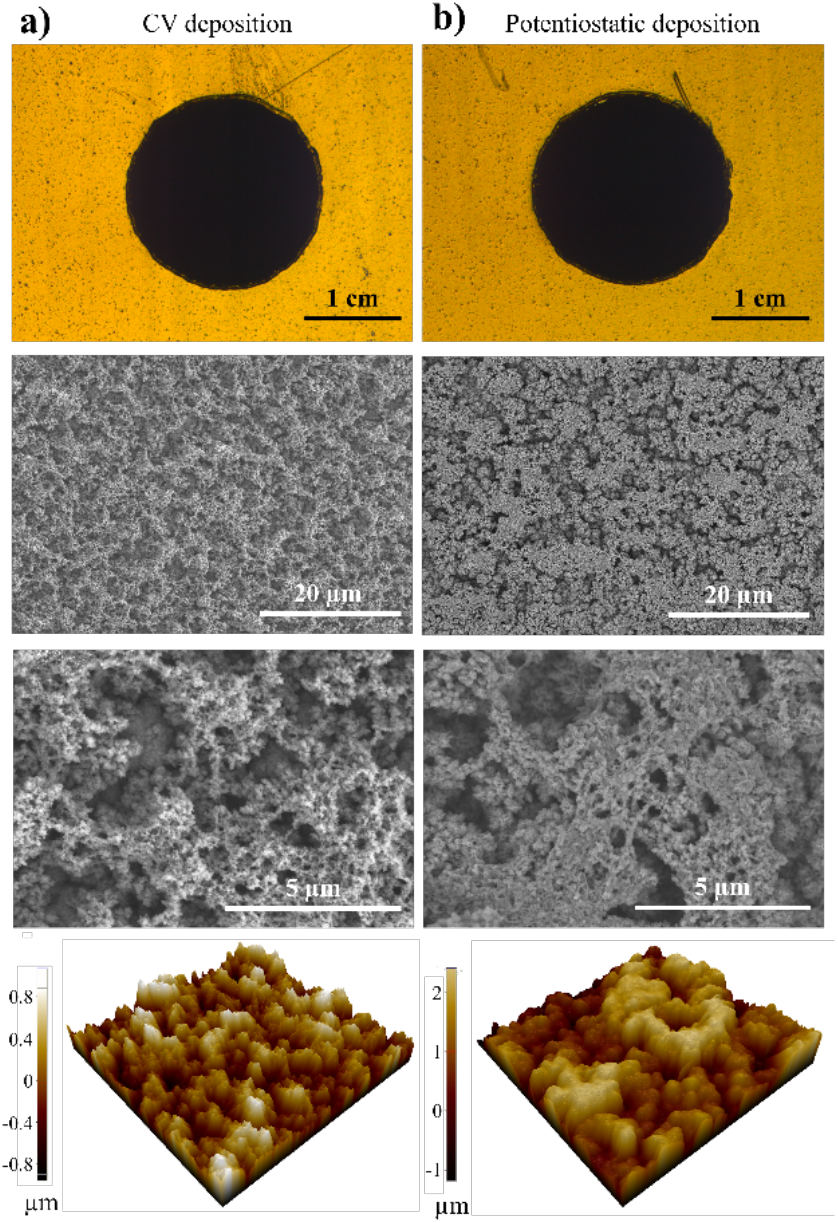
From top to bottom: Micrographs, SEM images; 25×25 µm^2^ AFM scan of the PEDOT:PDA films coated by a) CV-, and b) PS-deposition.

Under potentiostatic conditions, continuous nucleation and rapid growth lead to larger aggregates forming quickly, resulting in a more heterogeneous surface with coarser features. In contrast, CV employs multiple oxidation-reduction cycles that enable more controlled nucleation and gradual material addition. Each cycle builds upon the previous layer, producing a more uniform and finer-grained structure. Despite the morphological differences, both methods had comparable electrochemical performance, confirming overall similar EASA and ion accessibility. Films deposited by PS techniques were 45% thicker than with the CV technique, a difference explained by reversible redox cycling of dopamine during the cathodic sweeps in CV, where partially reduced leucodopamine redissolves, resulting in a net lower charge-to-thickness conversion compared to PS deposition. Overall, the increase in roughness and therefore EASA, coupled with PEDOT’s mixed electronic-ionic conduction, explains the enhanced performance regardless of the deposition technique.

### E. Structural analysis

FTIR spectroscopy confirmed the chemical composition of the PEDOT:PDA composite. The spectrum (Fig. 6) displays characteristic absorption bands of both PEDOT and PDA, confirming successful co-deposition. In the pure PEDOT spectrum, the peak at 2919 cm^−1^ is attributed to C–H stretching in the ethylenedioxy ring, while the peak at 1345 cm^−1^ corresponds to C–C inter-ring stretching in the thiophene ring [12]. The C–O–C stretching vibrations of the ethylenedioxy group appear at 1171 and 1054 cm^−1^, and the C–S stretching is identified at 904 and 843 cm^−1^ [13], [14]. In the pure PDA spectrum, the broad band at 3389 cm^−1^ is characteristic of O–H and N–H stretching vibrations. The peaks at 1591 and 1544 cm^−1^ correspond to aromatic C=C stretching, while those at 1379 and 1248 cm^−1^ are assigned to C–O–H bending in the catechol group. The peak at 2944 cm^−1^ is attributed to C–H stretching in the aryl group [15].

**Figure 6:**
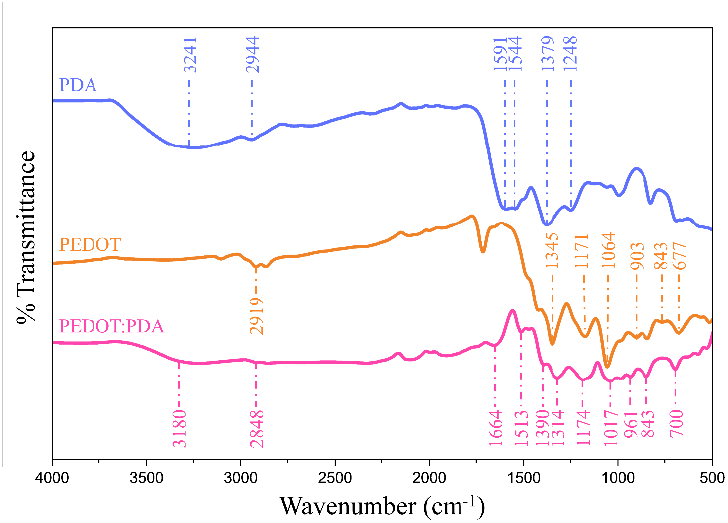
FTIR spectra of PEDOT, PDA and PEDOT:PDA coating

In the PEDOT:PDA composite, peaks from both components are identified. The broad O–H/N–H stretching band (~3200–3400 cm^−1^) in pure PDA is noticeably less intense in the PEDOT:PDA composite. The peak has shifted towards a lower wavenumber (3241 cm^−1^ in PDA compared to 3180 in PEDOT:PDA). This reduction indicates partial oxidation of catechol OH groups to quinones (corroborated by the new 1664 cm^−1^ C=O band) [16] and stronger hydrogen bond formation between PEDOT and PDA [17]. The C–H stretching band common to both PDA’s aryl group and PEDOT’s ethylenedioxy ring is observed at 2848 cm^−1^. The peak at 1513 cm^−1^ is characteristic of the C_α_=C_β_ of the PEDOT’s quinoid group [18]. The peak at 1390 cm^−1^ corresponds to phenolic C–O–H bending from PDA [19], and the peak at 1314 cm^−1^ to C–C inter-ring stretching from PEDOT [20]. The peaks at 1174 and 1017 cm^−1^ confirm the presence of PEDOT’s C–O–C stretching. Finally, the peaks at 961, 843 and 700 cm^−1^ matches those of the C-S stretching in the PEDOT. FTIR analysis confirms the formation of the PEDOT:PDA composite through several spectral changes. A new band at 1664 cm^−1^ appears exclusively in the composite spectrum, absent in both pure components. The 1513 cm^−1^ band, absent in PEDOT, appears in the composite. The C–C inter-ring stretching band shifts from 1345 cm^−1^ in pure PEDOT to 1314 cm^−1^ in the composite, reflecting changes in the polymer chain environment. Finally, the broad O–H/N–H stretching band of PDA (~ 3241 cm^−1^) decreases in intensity in the composite, suggesting chemical modification of PDA’s catechol groups upon interaction with PEDOT. Although these observations demonstrate interactions between PEDOT and PDA beyond simple physical mixing, further studies are needed to pinpoint precise molecular interactions.

## IV. Conclusion

In summary, we report a facile and scalable strategy for fabricating high-performance bioelectrodes via CV co-electropolymerization of PEDOT and polydopamine. The resulting hybrid films exhibit excellent electrochemical properties, and rough surfaces favorable for applications in implantable bioelectronics, sensors, and tissue-electrode interfaces. A comparison between potentiodynamic and potentiostatic depositions were done to investigate the differences in electrochemical performance and film morphology. Future work will aim at thoroughly characterize the full range of benefits of such hybrid biomaterial coatings such as cell-affinity and sensing capabilities.

## Data Availability

The dataset supporting this work is available at https://doi.org/10.5281/zenodo.20797822 [21].

